# Application of light-sheet mesoscopy to image host-pathogen interactions in intact organs

**DOI:** 10.1101/2022.04.04.486945

**Authors:** Eliana Battistella, Juan F. Quintana, Gail McConnell

**Author notes:** **Additional author contact information:**.

## Abstract

Human African Trypanosomiasis (HAT) is a disease caused by the extracellular parasite *Trypanosoma brucei* that affects the central nervous system (CNS) during the chronic stage of the infection, inducing neuroinflammation, coma, and death if left untreated. However, little is known about the structural change happening in the brain as result of the infection. So far, infection-induced neuroinflammation has been observed with conventional methods, such as immunohistochemistry, electron microscopy, and 2-photon microscopy only in small portions of the brain, which may not be representative of the disease. In this paper, we have used a newly-developed light-sheet illuminator to image the level of neuroinflammation in chronically infected mice and we use analyse these data to compare the infection in naïve controls. This system was developed for imaging in combination with the Mesolens objective lens, providing sub-cellular resolution for imaging volumes exceeding 39 mm^3^ in an acquisition time of only 8 hours. The mouse brain specimens were cleared using CUBIC+, followed by antibody staining to locate Glial Fibrillary Acid Protein (GFAP) expressing cells, primarily astrocytes and ependymocytes, used here as a proxy for cell reactivity and gliosis. The large capture volume allowed us to detect GFAP^+^ cells and spatially resolve the innate responses to *T. brucei* infection. Based on morphometric analyses and spatial distribution of GFAP^+^ cells, our data demonstrates a significant increase in cell dendrite branching around the lateral ventricle, as well as dorsal and ventral third ventricles, that are negatively correlated with the branch extension in distal sites from the circumventricular spaces. To our knowledge, this is the first report highlighting the potential of light-sheet mesoscopy to characterise the inflammatory responses of the mouse brain to parasitic infection at the cellular level in intact cleared organs, opening new avenues for the development of new mesoscale imaging techniques for the study of host-pathogen interactions.

## 1 Introduction

Chronic infections of the central nervous system (CNS), such as those caused by the protozoan parasite *Trypanosoma brucei*, causative agent of sleeping sickness, are typically associated with reactivity of glial cells (microgliosis and astrogliosis) and neuroinflammation (Lundkvist et al., 2004; Kristensson et al., 2010; Laperchia et al., 2016; Bentivoglio et al., 2018; Tesoriero et al., 2018; Rodgers et al., 2019a). However, limited information regarding the ultrastructural changes that take place in the CNS during sleeping sickness. During CNS colonisation, *T. brucei* is thought to exploit several routes, including the blood-brain barrier and the circumventricular organs to colonise the CNS (Kristensson et al., 2010), although the inflammatory responses in the brain parenchyma are poorly understood. Most of the studies to date employ methods such as conventional immunohistochemistry, electron microscopy, and more recently 2-photon microscopy, to observe infection-induced neuroinflammation (Laperchia et al., 2016, 2018; Rodgers et al., 2019b). However, these methods can only image small volumes of brain and thus little is known about the changes to whole brain architecture in response to infection.

Volumetric optical mesoscopic imaging, which bridges the gap between microscopy and macrophotography, offers a possible solution to overcome these technical limitations. There exist several mesoscale imaging technologies (Munck et al., 2021), but for studying whole mouse brain architecture with high spatial resolution in three dimensions the Mesolens is the ideal objective lens. It is a giant multi-element lens with high numerical aperture (0.47) and low magnification (4x) and achieves three-dimensional imaging with 700 nm lateral resolution and 4 µm axial resolution in a volume of capture exceeding 100 mm^3^ (McConnell et al., 2016). The acquisition of images with the Mesolens is already possible with brightfield illumination (Shaw et al., 2021), confocal laser scanning (McConnell et al., 2016) and widefield epifluorescence and HiLo modalities (Schniete et al., 2018). However, although the Mesolens confocal laser scanning modality has been successfully demonstrated with a variety of biological samples (McConnell and Amos, 2018; Francis et al., 2020; Rooney et al., 2020) acquiring a full resolution imaging volume is time consuming. The minimum acquisition time for a single image is 200 s (0.5 µs pixel dwell time for 20000 × 20000 pixels required for Nyquist sampling) resulting in a full volume acquisition time from a minimum of 55 hours to several weeks (McConnell et al., 2016). Light-Sheet Fluorescence Microscopy (LSFM) is now a routine tool in bioscience research and enables more rapid three-dimensional imaging than confocal laser scanning. The popularity of this technique is certainly due to the numerous configurations (Huisken et al., 2004; Tokunaga et al., 2007; Gao et al., 2012; Chen et al., 2014; Hoyer et al., 2016) that can be implemented to allow the investigation of a wide variety of specimens of many different sizes (Planchon et al., 2011; Tomer et al., 2012; Glaser et al., 2019; Voigt et al., 2019).

One of the many applications of LSFM is to explore the tissue architecture of complex organs, such as the CNS, in health and disease. For example, similar approaches have been employed to resolve structural features of neuronal synapsis to reconstruct activity patterns and connectivity maps (Niedworok et al., 2012; Bèchet et al., 2020; Blutke et al., 2020; Perens et al., 2021; Zhang et al., 2021). However, to our knowledge, LSFM has not been implemented to study infectious diseases affecting the CNS at the mesoscale. The recent development of a light-sheet illuminator for use in combination with the Mesolens has greatly increased the speed with which large volumetric imaging at the mesoscale could be obtained while preserving sub-cellular resolution (Battistella et al., 2022). However, this is not trivial from an optical engineering perspective. It requires either a long and thin static light-sheet through which the specimen is moved, or an accurate synchronisation between the camera detector and a swept light-sheet waist across the field of view. The swept light-sheet method is not possible with the Mesolens because of detection requirements (Battistella et al., 2022). We have developed and built two light-sheet illuminators for use with the Mesolens. The first uses a simple Gaussian beam light-sheet and is suitable for faster, low-resolution scans. The second is an Airy beam light-sheet, for higher spatial resolution imaging.

Here we have applied these light-sheets for rapid high-resolution mesoscale imaging and quantification of neuroinflammation during *T. brucei* infection using an experimental murine model of infection that faithfully recapitulates the severe neuroinflammatory state observed in patients infected with *T. b. rhodesiense* (Rodgers et al., 2015, 2019a; Lamour et al., 2017) observed a significant increase in the labelling intensity of the protein glia fibrillary acidic protein (GFAP) (Yang and Wang, 2015), used here as a proxy for astrocyte reactivity and inflammation, around the circumventricular organs (CVOs), coinciding with the proposed sites of entry of the parasites into the CNS (Bentivoglio et al., 2018). Concomitantly, GFAP positive cells (typically astrocytes and ependymocytes) located in the vicinity of the ventricular structures display a greater number of cytoplasmatic processes when compared to either parenchymal astrocytes in the infected brain, or those found in naïve controls. These data not only suggest a potential spatially resolved response to parasitic infections, driven by GFAP^+^ cells in the proximity to the CVOs, but also demonstrate the impact of light-sheet mesoscopy to resolve cellular responses to infection in intact organs.

## 2 Materials and methods

### 2.1 *Trypanosoma brucei brucei* strain

The *T. b. brucei* AnTat 1.1E clone is a pleomorphic strain that induce a chronic infection in mice. This strain was derived from a strain originally isolated from a bushbuck in Uganda in 1966 (D. Le Ray et al., 1977) and kept as pleomorphic lines for *in vivo* studies.

### 2.2 Experimental infections with *T. b. brucei* and brain clearing

All animal experiments were approved by the University of Glasgow Ethical Review Committee and performed in accordance with the home office guidelines, UK Animals (Scientific Procedures) Act, 1986 and EU directive 2010/63/EU. All experiments were conducted under SAPO regulations and project licence number: 200120043.

We used an experimental chronic infection system using the tolerant C57BL/6 mouse strain inoculated with *T. brucei brucei* Antat 1.1E, which is a well-characterised model in which parasite start colonising the CNS around 21 days post-infection (Lundkvist et al., 2004; Laperchia et al., 2016; Rijo-Ferreira et al., 2018). Eight-week-old female C57Black/6J mice (Jackson labs, UK) were inoculated by intra-peritoneal injection with 10^4^ *T. b. brucei* Antat 1.1E. Parasitaemia was monitored every two days from the superficial tail vein, examined using phase microscopy, and scored using the rapid “matching” method (Herbert and Lumsden, 1976). Uninfected mice of the same strain, sex and age served as uninfected controls. Following the parasite infection, we noted two main peaks of parasitaemia around day 4 and day 14 post-infection (Figure 2E). At 21 days post-infection, mice were culled by cervical dislocation under terminal anaesthesia with isofluorane and decapitated to remove intact brain. Using absorption staining and brightfield microscopy we could readily detect parasites in the brain parenchyma at this point of infection and compare it with the naïve control (Figure 2A-D), which is consistent with previous results at the microscopic scale (Laperchia et al., 2016).

After an overnight fixation in 10% neutral buffered formalin, samples were cleared using CUBIC (TCI Chemicals) (Tainaka et al., 2018). Briefly, samples were first delipidated using CUBIC-L for a period of ∼6-7 days at 37 °C. After clearing was completed, samples were washed three times in 1X PBS (including an overnight wash at 4 °C), blocked for 2 hours at room temperature with 1X PBS supplemented with 5% Fetal Calf Serum (Invitrogen) and 0.25% Tween-20 (Invitrogen), and stained with anti-GFAP mouse monoclonal antibody coupled to Alexa Fluor 488 (Clone GA5, 1/100, Thermo) for 3 days at 37 °C and gentle agitation. We chose this protein, mainly expressed by astrocytes and to a lesser extent by ependymocytes (Roessmann et al., 1980), as a target for imaging experiments because the expression level of this protein is traditionally used as a proxy to examine astrogliosis, including those induced during sleeping sickness (Yang and Wang, 2015). Samples were then washed three times with 1X PBS to remove excess antibody and incubated in CUBIC-R+ (TCI chemicals) for 3 days for refractive index matching.

### 2.3 Histological and immunohistochemistry evaluation of coronal brain sections

Paraformaldehyde-fixed coronal brain sections were trimmed and processed into paraffin blocks. 3 µm thick sections were processed for immunohistochemical staining using a polyclonal rabbit antibody raised against the *T. brucei* Heat Shock Protein 70 (TbHSP70) (Kind gift from Professor Keith Matthews, Edinburgh, UK) using a Dako Autostainer Link 48 (Dako, Denmark) and were subsequently counterstained with Gill’s haematoxylin.

### 2.4 Development of light-sheet illuminators for mesoscale imaging

We developed two different light-sheets for use with the Mesolens. The first was a simple and long but relatively thick light-sheet using Gaussian optics. The second was a propagation-invariant light-sheet using an Airy beam, which covers the same area as the Gaussian light-sheet but is a fraction of the thickness for improved optical sectioning. A Coherent Sapphire 488-10 CDRH laser emitting light at 488 nm with a maximum of 10 mW average power at the sample plane was used as light source used for both light-sheet illuminators. The Gaussian light-sheet was formed by directing the 8 mm expanded beam through a cylindrical lens (f = 300 mm. LJ1558RM-A, Thorlabs). The Airy light-sheet was shaped with the following optical elements: two aspherical lens (AL, f = 20mm. AL2520-A, Thorlabs), a Powell lens (PL, fan angle = 10°, Laserline optics Canada), two cylindrical lenses (CL, f = 75mm. LJ1703RM-A, Thorlabs). The schematic of both setups is shown in Figure 1. A more detailed description of the optical arrangement and characterisation of the light-sheet imaging performance is available (Battistella et al., 2022).

**Figure 1.**
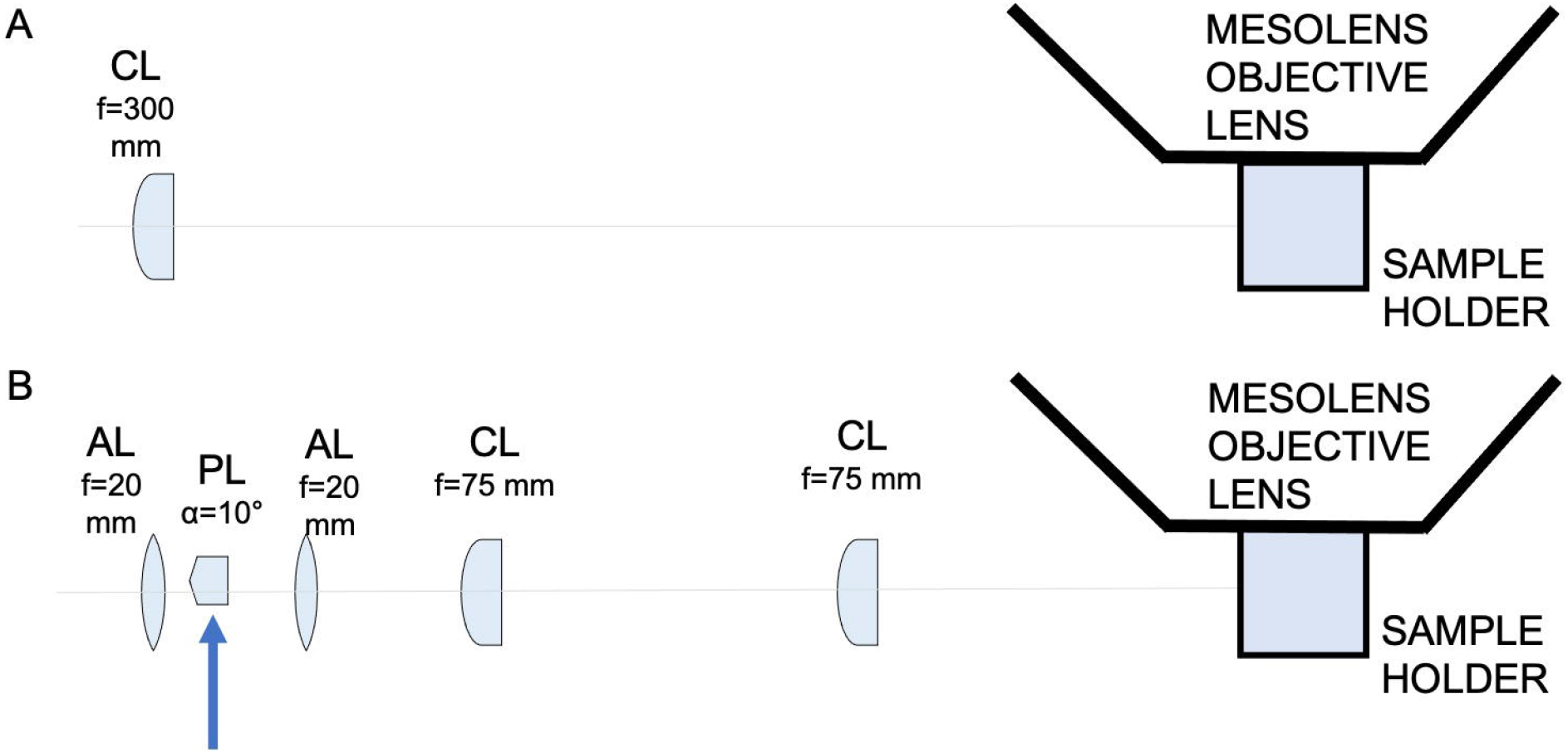
Schematic diagram showing the light-sheet illuminators. (A) Schematic diagram showing the Gaussian light-sheet illuminator. The light-sheet is shaped using a cylindrical lens (CL, f = 300 mm) and directed to the sample plane. The sample is placed in a quartz cuvette to be imaged by the Mesolens. (B) Schematic diagram showing the Airy light-sheet illuminator. The light-sheet is shaped using a combination of aspherical lenses, aspheric lens (AL, f = 20 mm), a Powell lens (PL, fan angle = 10°), aspheric lens (AL, f = 20mm), two identical cylindrical lenses (CL, f = 75 mm). The Powell lens was moved out of the optical axis to create the Airy light-sheet, as indicated by the blue arrow in the schematic.

### 2.5 Imaging of mouse coronal brain sections

The cleared brain specimens were mounted in CUBIC+ mounting media in a quartz cuvette (CV10Q3500F, Thorlabs). The open end of the cuvette was closed with a 3D-printed stopper and sealed with clear nail varnish. A 60 mm diameter borosilicate coverslip (0107999098, Marienfeld) was placed on top of the cuvette to support oil immersion with type LDF immersion oil (16241, Cargille). The Mesolens objective is equipped with correction collars to minimise spherical aberrations and in this work the collars were set for to match the refractive indices of CUBIC+ and the LDF oil. The specimen was oriented so that an optical section corresponded to a coronal section, with the retrosplenial area being the first part of the specimen illuminated by the light-sheet. The region considered in the first instance was visibly identified by the ventral part of the retrosplenial area and by the part of the corpus callosum. The specimen was then translated along the y direction to image both the thalamus and hypothalamus in the infected brain specimen. The axial displacement z = 1.5 µm for the Airy light-sheet illumination was chosen to satisfy the Nyquist sampling criterion (Shannon, 1949), while z = 3.0 µm was used for the Gaussian light-sheet illumination.

The image datasets were acquired using the custom ‘MesoCam’ software, which controlled a sensor-shifting camera and acquired images of 12-bit depth in the TIFF format (Schniete et al., 2018). Nyquist sampling in the lateral dimension for the Mesolens FOV in widefield (and therefore light-sheet) modality corresponds to a 224 nm pixel size, for a total of 260 MP. There are no commercially available cameras with said resolution, but the Mesolens camera detector (VNP-29MC, Vieworks) is equipped with a chip-shifting mechanism that by shifting the image by 1/3 of a pixel in a 3×3 square allows Nyquist sampling, as described in a previous publication (Schniete et al., 2018). All images were acquired with 1000 ms exposure time for each of the 9 sensor positions and with a gain setting equal to 50. The acquisition of a single optical section took 9 seconds; however, the overall acquisition speed was limited by the time employed by the MesoCam software to transfer and save the acquired image from the camera to the PC. The acquisition volume for both naïve and infected specimens in the Airy light-sheet modality was composed of 933 slices, at the full 3.0 mm x 4.4 mm FOV. The entire volume was acquired in approximately 8 hours.

### 2.6 Image post-processing and visualisation

The images were visualised in two-dimensions and processed with Fiji (Schindelin et al., 2012) using virtual stacks. The outliers due to camera noise were removed using the batch processor for virtual stacks with the ‘RemoveOutliers’ function with a radius equal to 3.

A maximum intensity projection for each dataset was computed to allow for a qualitative evaluation of the datasets through two-dimensional visualisation. This was achieved with the ‘Z Project’ function of Fiji. The three-dimensional data handling and visualisation was possible thanks to a Python script using the Dask library (Dask Development Team, 2016) and the visualisation framework napari (napari contributors, 2019). Imaris (Oxford Instruments 2021) was used for the three-dimensional visualisation, rendering, and the fluorescence intensity calculations. We measured the fluorescence intensity to quantify the presence or the absence of GFAP+-reactive cells in each specimen, as the astrocytes reactivity and inflammation can indicate the sites of entry of the parasites into the CNS (Bentivoglio et al., 2018). To calculate the fluorescence intensity in both the naïve and in the infected specimens, the total fluorescence intensity level was measured from each volume using the Data Intensity variables of Imaris.

As Airy light-sheet illumination is not a widely used modality it was essential to evaluate whether the use of deconvolution algorithm is a valid method to improve the datasets resolution and contrast without introducing artefacts. Deconvolution was achieved using the Huygens software (Scientific Volume Imaging). The deconvolution was performed on a sub-volume composed of 100 optical slices, which corresponds to a total depth of 150 µm and an approximately 50 GB file size. This particular sub-volume file size was chosen since our computational power was limited by the available server RAM. The process was time consuming, exceeding 12 hours computational time per dataset. For the Airy light-sheet deconvolution parameters were chosen to approximate a Gaussian light-sheet with a 12 µm diameter beam waist, directed from the bottom of the image, and with no focus offset. The other parameters were set to describe the excitation and peak emission wavelengths of green fluorescent protein (488 nm and 520 nm), the sampling intervals (x and y: 224.4 nm, z: 1500 nm), and the optical parameters (numerical aperture 0.46, LDF oil immersion refractive index: 1.515).

Image post-processing was performed subsequently to the image acquisition and therefore did not affect the acquisition time. The removal of outliers took approximately 3 hours, and the maximum intensity projection was performed in a few minutes. Deconvolution of the selected sub-volumes were performed overnight.

### 2.7 Dendrite length measurement

The volume comprising the right ventricle was sampled with 35 cubic sub-volumes of 200 µm side, positioned at the same sample depth (approximately midway in the acquisition volume). The sub-volumes were chosen to be at a fixed distance from the separation of the ventricles (200 µm, 400 µm, 600 µm, 800 µm, 1000 µm, 1400 µm, 1800 µm) and distributed along 5 lines, separated by a distance of 200 µm. The dimension of the sub-volumes was chosen to guarantee the presence of at least 10 astrocytes. For each sub-volume the dendrite of 10 astrocytes were manually tracked using the AutoPath method for Filaments in Imaris. When a sub-volume contained more than the previously defined number of astrocytes, 10 astrocytes were randomly selected. The mean dendrite length, its standard deviation and the dendrites count for the tracked astrocytes was evaluated for each sub-volume using “Dendrite Length” information in the Imaris Detailed Statistics tab. The standard deviation and mean for same-distance sub-volumes were then combined to form a single group. The same method was used for the left ventricle.

## 3 Results

### 3.1 Airy light-sheet imaging offers improvement in resolution over Gaussian light-sheet mesoscopy

Gaussian illumination is a standard in light-sheet microscopy therefore a comparison with our newly developed Airy illuminator was essential to evaluate the optical sectioning capability and the imaging performances of the two light-sheet modalities. The same infected mouse brain specimen was imaged at the same volume depth by rotating the cuvette by 90° so that the Gaussian light-sheet travels through the same clear side of the cuvette as for the Airy light-sheet illumination. A comparison between two single optical sections, acquired with the two different light-sheet modalities is shown in Figure 3A and 3B. The digitally zoomed regions of interest shown in Figure 3C, and Figure 3D confirms the sub-cellular resolution detail possible with the Airy light-sheet, where dendrites and filaments are observed, compared with the Gaussian illuminator, which only just resolves the dendrites. Moreover, the comparatively thick Gaussian light-sheet reveals autofluorescence from cell bodies, potentially originating from poor delipidation in the observed area. This cellular autofluorescence is not present in the Airy light-sheet images, thus facilitating a more straightforward quantitative analysis of image datasets. Notably, these images also included vertical stripes which are characteristic light-sheet artefacts that are generated by the sample scattering and absorption (Stainier and Huisken, 2007). We also noted that the stripe artefacts in the images acquired with the Gaussian beam light-sheet are approximately 4-times brighter than those compared to those obtained using the Airy light-sheet illuminator. Figure 3E shows the relative percentage comparison of the fluorescence intensity region with stripe artefacts in Figure 3A and 3B. This 4-fold increase may be a consequence of the increased light-sheet thickness for the Gaussian beam.

**Figure 2.**
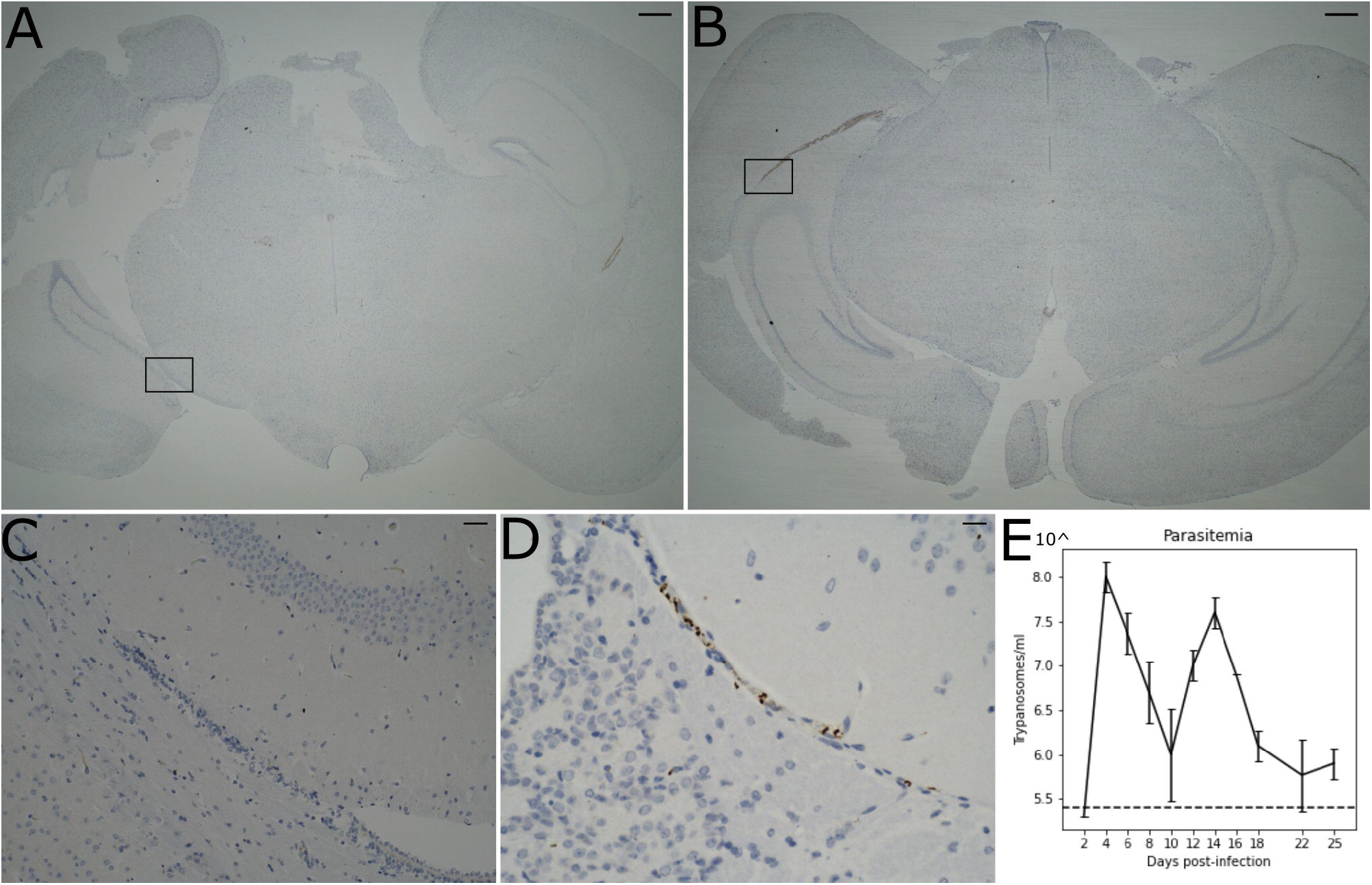
Presence of parasite in the brain parenchyma. (A) Histological brightfield images highlight the presence of parasites in the brain parenchyma in the naïve and (B) infected specimens. Scale bar = 100 µm The boxed region in (A) and (B) are magnified in (C) e (D). Scale bar = 25 µm. (E) Level of parasitaemia in the infected mouse averaged over three biological replicates in a time course of 25 days. The dotted line represents the detection limit of the instrument (10^5.4^ trypanosomes/ml).

**Figure 3.**
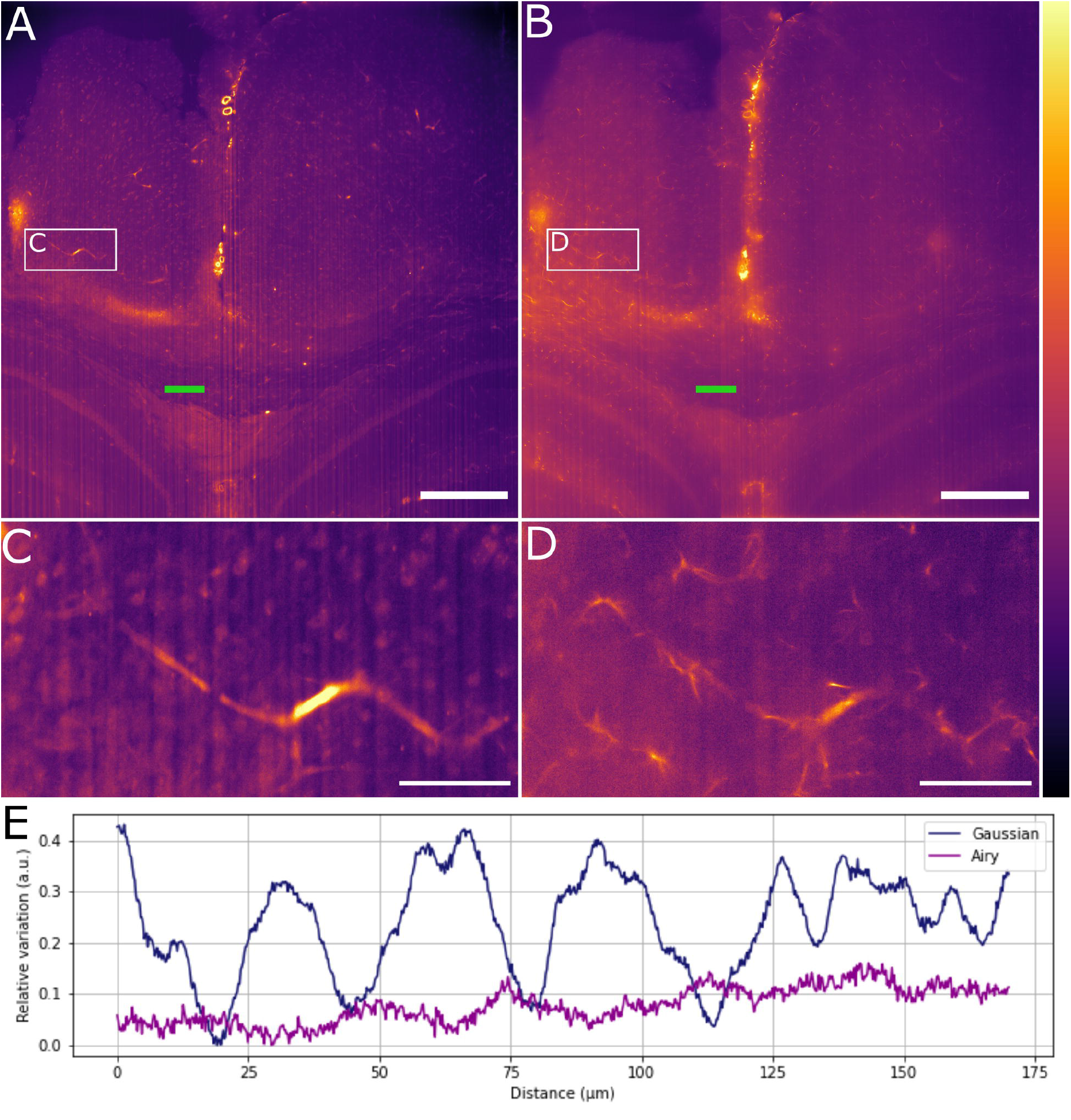
Comparison between Gaussian light-sheet and Airy light-sheet images. (A) Single optical section of the same infected mouse brain, acquired with the Gaussian light-sheet and (B) the Airy light-sheet setup. The boxed region in (A) and (B) are magnified in (C) e (D) to highlight the increase in subcellular details obtained with the Airy light-sheet. The look-up table for (A)-(D) is mpl-inferno. The scale bar is respectively (A)-(B) 500 µm and (C)-(D) 100 µm. The images were converted to 8-bit values (0-255). (E) Comparison between the relative variation of the stripes artefact intensity in the central region of the specimen indicated by the green line in panels A and B.

### 3.2 Deconvolution further improves Airy light-sheet imaging of inflammation

We have shown how the use of the Airy light-sheet illuminator improves the imaging resolution. Deconvolution is known to improve the resolution of imaging systems but to our knowledge there is no report on how the proprietary deconvolution software Huygens (Scientific Volume Imaging) performs on imaging data obtained with non-Gaussian light-sheet illumination. We tested its performance on a sub-volume of the dataset shown in Figure 3A with the intent of exploring the potential of the software with this specific specimen type and imaging system. As the software does not provide an option for static Airy light-sheet illuminators, we aimed to understand whether processing our dataset using deconvolution methods would result in artefacts being created in the image caused by the presence of subsidiary peaks. Therefore, a 150 µm-thick sub-dataset was processed with the Huygens deconvolution software and the MIP is shown in Figure 4A. As expected, a consequence of the deconvolution process was an increase in image contrast, while the out-of-focus noise and background blur were reduced. By a visual analysis we could not detect any artefacts emerging from the deconvolution process. This would, in principle, facilitate the use of a segmentation algorithm if necessary for quantitative analysis. The presence of the astrocytes and ependymal cells is highlighted in different regions across the specimen (Figure 4B-D).

**Figure 4.**
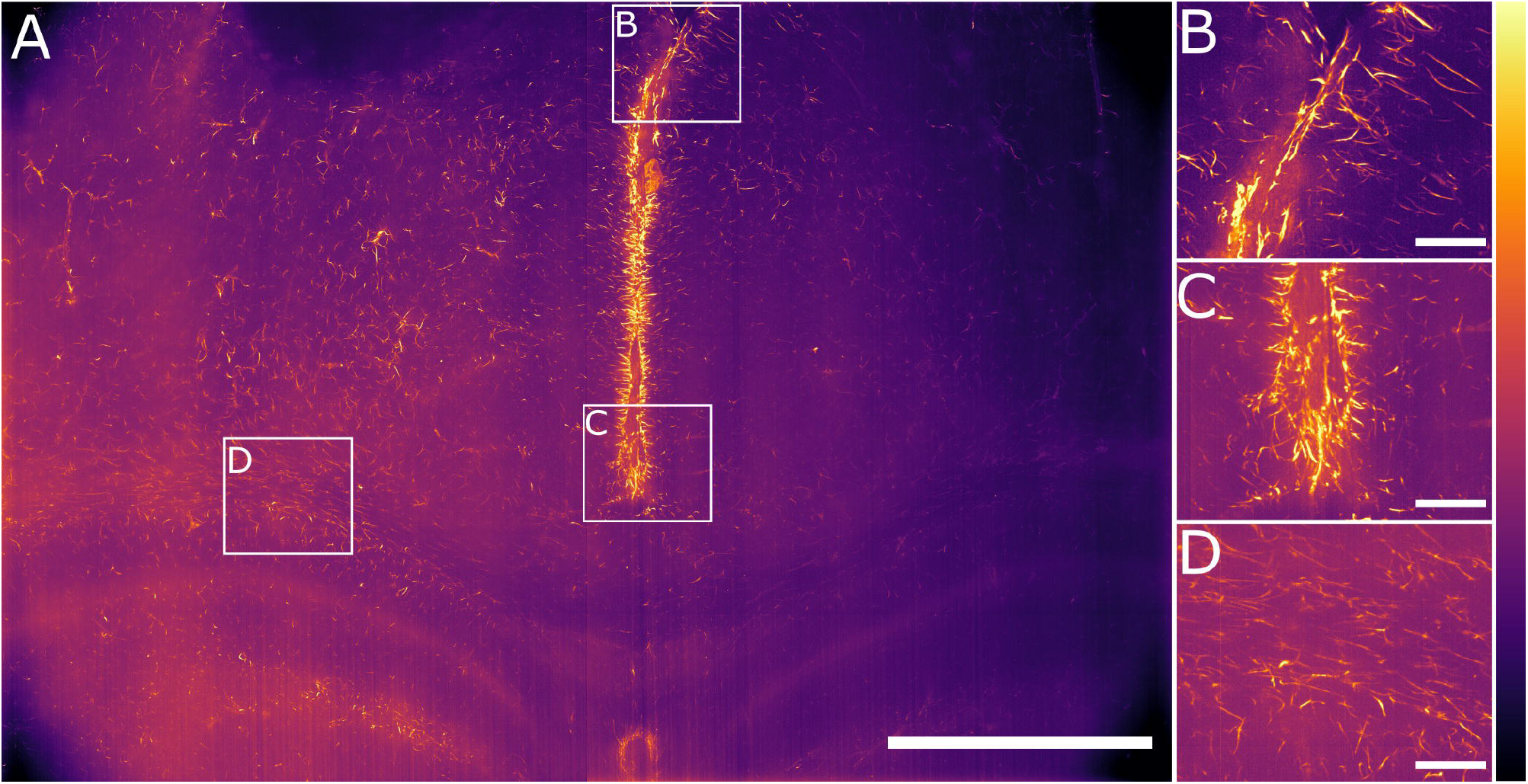
Maximum intensity projection of the sub-volume shown in Figure 3B after deconvolution. The full FOV is shown in (A). (B)-(C) highlight sub-regions in which both astrocytes and ependymal cells are resolved. (D) focuses on a sub-region in which the sole presence of GFAP reactive astrocytes is detected. The look-up table is mpl-inferno. The scale bar is respectively (A)-(B) 500 µm and (C)-(D) 100 µm. The images were converted to 8-bit values (0-255).

### 3.3 Airy light-sheet imaging of the chronically infected mouse brain shows GFAP+ reactivity in the vicinity of the circumventricular organs

We globally assessed the fluorescence intensity in different brain regions acquired with the Airy light-sheet setup from both naïve and the infected specimens. Overall, we observed an increased GFAP^+^ reactivity in all the areas evaluated, including the cingulate cortex (Figure 5A and 5B), corpus callosum and external capsule (Figure 5C and 5D), and hypothalamic area, close to the fornix and the median forebrain bundle (Figure 5E and 5F). Interestingly, these brain regions are typically associated with sites of parasite accumulation such as CVOs and leptomeninges. The relative difference in reactive GFAP+ cells in both the naïve and infected specimens was quantified by comparing the sum of the fluorescence intensity signal in both the whole field of view, as well as in selected sub-regions. A three-dimensional rendering of the regions in Figure 5A and 5B are shown in Animation1 and Animation2. The reported values are the sum of the pixel intensity for the full volumes of interest, knowing that each pixel in the acquired volume can display a value between 0 and 4095 (the camera pixel depth is 12-bit). We observed a total of 2.2. 10^13^ fluorescent counts in the naïve specimen compared to 3.0. 10^13^ fluorescent counts in the infected specimen when assessing the cingulate cortex area indicating an increase of 38% in GFAP^+^ reactivity upon infection. Subsequent region-specific measurements suggest a localised increased GFAP^+^ reactivity of 52% in the corpus callosum and external capsule area and 103% in the hypothalamic area. The values are summarised in Table 1. The comparison between the naïve and the infected specimens in Figure 5 shows that the fluorescence signal of the naïve specimen can be attributed mostly to background signal possibly constituted by poor delipidation, non-specific staining, and basal expression of GFAP in astrocytes and ependymal cells (Roessmann et al., 1980). Therefore, the fluorescence increase measured for each naïve-infected pair is attributed to fluorescence from GFAP^+^ cells, likely to be reactive astrocytes (Roessmann et al., 1980).

**Table 1:** Fluorescence intensity values for the region of the specimen shown in Figure 5.

|  | Naïve specimen<br>Intensity (counts) | Infected specimen<br>Intensity (counts) | Fluorescence intensity increase<br>((infected-naïve)/naïve) |
| --- | --- | --- | --- |
| Cingulate cortex<br>(Figures 5A – 5B) | $2.19 \cdot 10^{13}$ | $3.02 \cdot 10^{13}$ | 38% |
| Corpus callosum<br>and external capsule<br>(Figures 5C – 5D) | $1.03 \cdot 10^{13}$ | $1.57 \cdot 10^{13}$ | 52% |
| Hypothalamic area<br>(Figures 5E – 5F) | $0.60 \cdot 10^{13}$ | $1.22 \cdot 10^{13}$ | 103 % |

**Figure 5.**
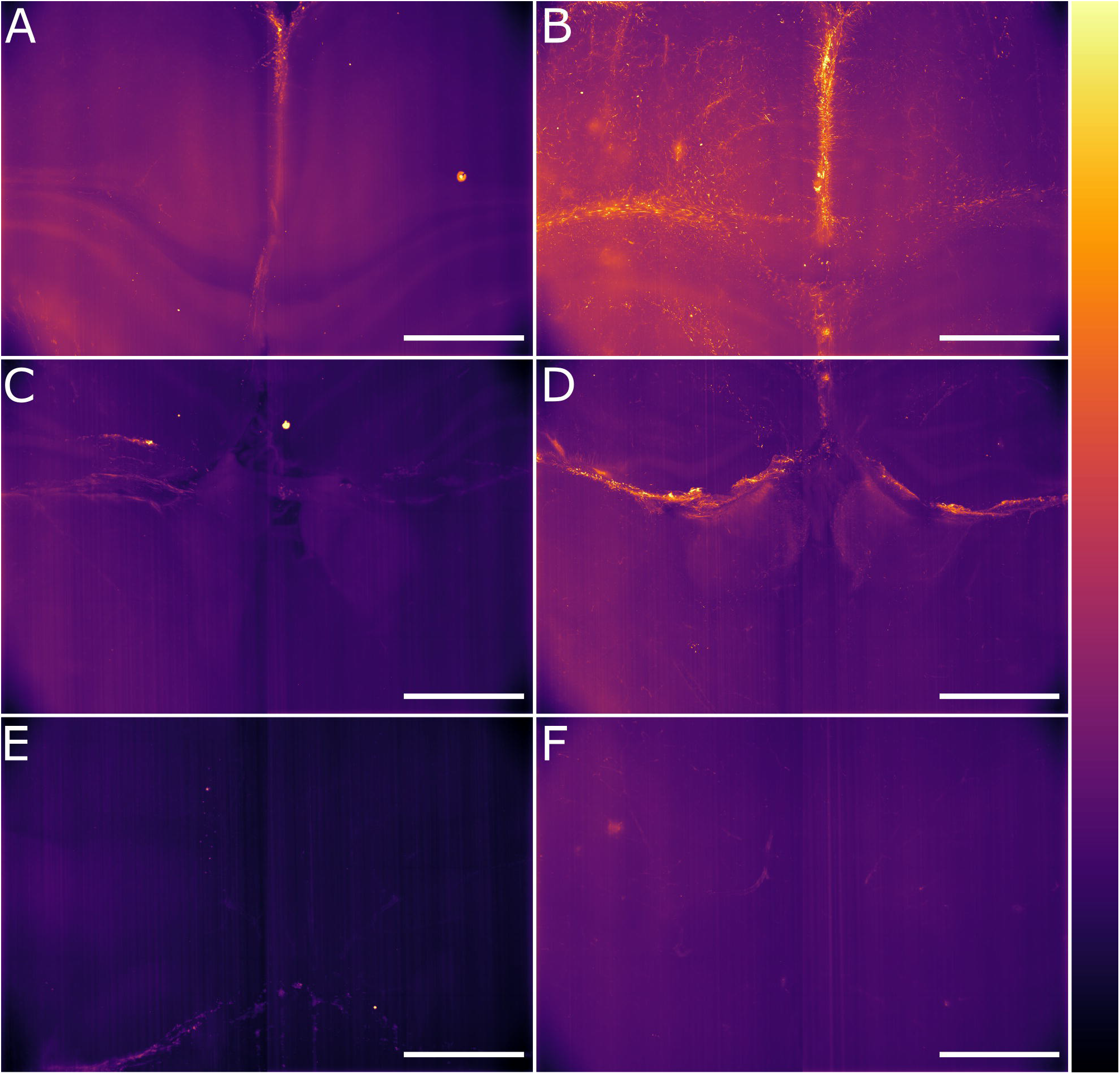
Light-sheet Mesolens images of comparable volumes in the naïve and infected specimens. We imaged and processed images from three main brain regions: cingulate cortex in the (A) naïve and (B) infected specimens, corpus callosum and external capsule in the (C) naïve and (D) infected specimens, and hypothalamic area, including the fornix and the forebrain bundle in the (E) naïve and (F) infected specimens. The look-up table is mpl-inferno. Scale bar: 1000 µm. The images were converted to 8-bit values (0-255).

### 3.4 Dendrite length evaluation reveals spatially resolved response to neuroinflammation

Having established a pipeline for acquiring and deconvolving the samples obtained from the light-sheet Mesolens, we performed a morphometric analysis of the different GFAP^+^ cell types detected in the specimens including in this study. More specifically, we measured dendrite length as a proxy for the typical morphological changes, including filopodia projections, observed in reactive astrocytes (Bailey and Shipley, 1993; Hochstim et al., 2008; Schiweck et al., 2018; Hyvärinen et al., 2019). Given the previous regional diversity of GFAP reactivity surrounding structures in proximity to the CVOs, we measured the average dendrite length at different distances, starting from the ventricular spaces (higher GFAP reactivity). The average dendrite length in the left ventricle showed a decreasing trend, starting from 26.1 µm ± 13.1 µm at 200 µm distance down to 17.4 µm ± 10.8 µm at 1000 µm distance and then increasing again to 21.3 µm ± 13.3 µm, 1800 µm away from the ventricular space and the circumventricular organs (CVOs). A diagram of this region is shown in Figure 6A, while Figure 6B shows a sub-region in which astrocytes were manually tracked. The dendrites in the right ventricle followed a similar and significant trend, showing a decrease from 26.0 µm ± 13.6 µm at 200 µm distance down to 18.6 µm ± 11.1 µm at 800 µm distance and then increasing again to 20.7 µm ± 12.4 µm, at 1400 µm to decrease again to 20.1 µm ± 10.8 µm,1800 µm away from the ventricle separation. These measurements are summarised in Table 2A and 2B. Of note, the same results were observed in both ventricles, suggesting that the inflammatory responses are bilateral in the brain.

**Table 2A:** Average dendrite length in the right ventricle measured at increasing distances from the circumventricular organs (CVOs). The p-value is less than 0.0005 for each sub-volume pair.

| Distance from the<br>CVOs ($\mu\text{m}$) | Mean length ($\mu\text{m}$) | Standard deviation ($\mu\text{m}$) | p-value |
| --- | --- | --- | --- |
| 200 | 26.0 | 13.6 | / |
| 400 | 20.6 | 12.2 | $4.3 \cdot 10^{-5}$ |
| 600 | 19.4 | 12.5 | $1.1 \cdot 10^{-6}$ |
| 800 | 18.6 | 11.1 | $4.1 \cdot 10^{-9}$ |
| 1000 | 19.0 | 10.8 | $3.7 \cdot 10^{-8}$ |
| 1400 | 20.7 | 12.4 | $1.2 \cdot 10^{-4}$ |
| 1800 | 20.1 | 10.8 | $6.2 \cdot 10^{-6}$ |

**Table 2B:** Average dendrite length in the left ventricle measured at increasing distances from the circumventricular organs (CVOs). The p-value is less than 0.0005 for each sub-volume pair.

| Distance from the<br>CVOs ($\mu\text{m}$) | Mean length ($\mu\text{m}$) | Standard deviation ($\mu\text{m}$) | p-value |
| --- | --- | --- | --- |
| 200 | 26.1 | 13.1 | / |
| 400 | 21.5 | 11.7 | $5.2 \cdot 10^{-4}$ |
| 600 | 19.5 | 12.9 | $3.4 \cdot 10^{-2}$ |
| 800 | 18.7 | 11.7 | $3.4 \cdot 10^{-9}$ |
| 1000 | 17.4 | 10.8 | $7.7 \cdot 10^{-14}$ |
| 1400 | 18.8 | 12.3 | $8.7 \cdot 10^{-9}$ |
| 1800 | 21.3 | 13.3 | $5.0 \cdot 10^{-4}$ |

**Figure 6.**
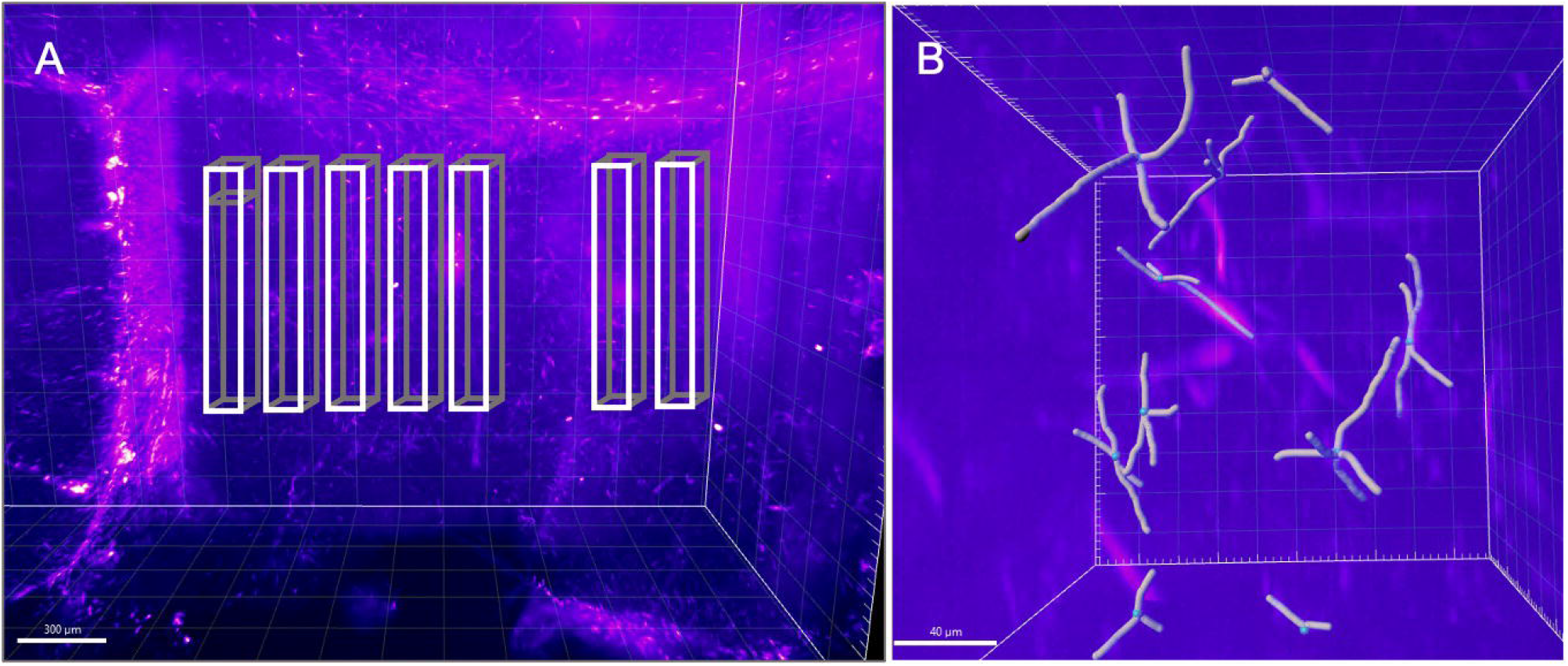
Schematic of the areas in which the astrocytes were tracked. (A) Area of the right ventricle in which the astrocytes were manually tracked. Each box was divided in 5 sub-boxes of 200 µm^3^ for ease of tracking. (B) Example of the sub-region in which 10 astrocytes were manually tracked. The scale bar is respectively (A) 300 µm and (B) 40 µm.

Taken together, these morphometric analyses reinforced our previous observations demonstrating a localised astrocytic response, with increased astrocyte hypertrophy in regions in close proximity to the CVOs.

## 4 Discussion

Chronic CNS infection with African trypanosomes leads to a myriad of neuropsychiatric disorders which are far from being fully understood. Here, we set out to develop and implement novel imaging methods, combining light-sheet and optical mesoscopy, to study the innate responses to infections with *T. brucei* in the intact mouse brain. One key observation is the spatially resolved inflammatory response in proximity to the circumventricular organs (CVOs), including the cingulate cortex and the corpus callosum, as measured by the levels of GFAP reactivity and astrocyte hypertrophy. In the cingulate cortex, the infected brain displays an increase in the fluorescence signal of 38% which directly relates to the presence of GFAP^+^ reactive astrocytes. An increased GFAP^+^ reactivity of 52% was measured also in the corpus callosum and external capsule area and of 103% in the hypothalamic area. Although especially in the hypothalamic area the light-sheet quality deteriorates due to the long optical path travelled inside the specimens and it is no longer possible to observe the single astrocytes, it was possible to estimate the increased fluorescence signal.

In addition to the overall GFAP fluorescence intensity, we also report a change in the average dendrite length at different distances from the ventricle separation. This gave us an estimate of the neuroinflammatory response following *T. brucei* infection. The tracked cells displayed a decreasing trend in the dendrite length, which followed the distance from the ventricles division. We observed a similar trend for both ventricles, indicating a symmetrical response to the infection. As the astrocytes were manually tracked, only a total sub-volume of 0.28 mm^3^ and 350 cells were considered in both the left and the right ventricles. However, with automatic three-dimensional segmentation a more extensive evaluation can be performed, including a larger number of cells.

We have shown the advantage of using an Airy light-sheet in combination with the Mesolens. Although the thicker Gaussian light-sheet is useful for imaging larger structures such as blood vessels, capillaries, and cell nuclei, the comparison of the Gaussian and the Airy light-sheet setups demonstrated that the thinner Airy light-sheet was essential to reveal the presence of GFAP^+^ reactive astrocytes. However, the Gaussian light-sheet illuminator is a good alternative capable of a faster acquisition when a high-resolution image is not required. Depending on the structures of interest, either Gaussian or Airy light-sheet modes can provide the user with volumetric information. In both cases the acquisition speed is mainly limited by the camera and software data handling, as discussed in detail in previous work regarding Mesolens widefield acquisition (Schniete et al., 2018; Battistella et al., 2022). Nevertheless, light-sheet acquisition with the Mesolens in both Gaussian and Airy modalities is significantly faster than for Mesolens confocal laser scanning as the same imaging volume would require several hundred hours for completion.

We have evaluated that, if deconvolution is required, the use of proprietary deconvolution software gives good results when using standard Gaussian parameters in first approximation. The deconvolution settings do not consider the Airy nature of the light-sheet, but we did not detect any imaging artefacts in the deconvolved images. Instead, the use of the deconvolution increased the image contrast and details resolution compared to the non-deconvolved data. This can allow, in combination with three-dimensional segmentation methods, a more accurate analysis of the astrocyte number and location. However, although the Airy light-sheet theoretically has the properties of being non-diffractive and self-healing, the quality of the illumination slightly deteriorates with the specimen penetration. As such, it is essential to have well-cleared specimens for deep brain imaging. We also note that the stripe artefacts present in our image datasets did not have an appreciable effect on our analysis.

In future work, the acquisition volume can be increased by approximately 1 mm by customising a cuvette enclose with an open top, sealed only by a glass coverslip. This will be advantageous for imaging a variety of larger specimens, noting also that this light-sheet can be compatible with different clearing and staining methods.

## Supporting information

Animation 1

Animation 2

## 7 Data Availability Statement

All data underpinning this publication are publicly available on request from the University of Strathclyde KnowledgeBase at: https://doi.org/10.15129/4c1c6de5-3e1c-483d-a95b-e957dfad4e9f

## 9 Author Contributions

Light-sheet development, imaging, image processing, data analysis EB; infection, clearing, histology JFQ; experiments design and manuscript writing EB, JFQ, GM.

